# Close to optimal cell sensing ensures the robustness of tissue differentiation process: the avian photoreceptor mosaic case

**DOI:** 10.1101/2021.02.14.431147

**Authors:** Arnab Barua, Alireza Beygi, Haralampos Hatzikirou

## Abstract

The way that progenitor cell fate decisions and the associated environmental sensing are regulated to ensure the robustness of the spatial and temporal order in which cells are generated towards a fully differentiating tissue still remains elusive. Here, we investigate how cells regulate their sensing intensity and radius to guarantee the required thermodynamic robustness of a differentiated tissue. In particular, we are interested in finding the conditions where dedifferentiation at cell level is possible (microscopic reversibility) but tissue maintains its spatial order and differentiation integrity (macroscopic irreversibility). In order to tackle this, we exploit the recently postulated Least microEnvironmental Uncertainty Principle (LEUP) to develop a theory of stochastic thermodynamics for cell differentiation. To assess the predictive and explanatory power of our theory, we challenge it against the avian photoreceptor mosaic data. By calibrating a single parameter, the LEUP can predict the cone color spatial distribution in the avian retina and, at the same time, suggest that such a spatial pattern is associated with quasi-optimal cell sensing. By means of the stochastic thermodynamics formalism, we find out that thermodynamic robustness of differentiated tissues depends on cell metabolism and cell sensing properties. In turn, we calculate the limits of the cell sensing radius that ensure the robustness of differentiated tissue spatial order. Finally, we further constrain our model predictions to the avian photoreceptor mosaic.

## 1. Introduction

Decision-making is a process to identify important choices and responses which depends on some basic criteria [1]. Cell decision-making is a process where cells select a new state, such as cell fates or phenotypes, in response to their microenvironmental milieu. In this regard, pluripotent cell differentiation can be viewed as a cell decision-making of inheritable fates. Typically, cells irreversibly acquire new fates by following a hierarchical lineage, where pluripotent stem cells in a proper microenvironment (stem cell niche) differentiate into, for example, bone, muscle, epithelial, and further specialized cells. Cell differentiation process encompasses the dramatic change of geometry, shape, gene expression inside the cell, etc. [2,3]. It is yet to be fully understood how the information available to pluripotent progenitors, including its intrinsically determined state and extrinsic microenvironmental signals, is encoded and processed by progenitors to generate different differentiated cell types.

In 2006, Takahashi and Yamanaka [4] discovered that almost any differentiated cell can be sent back in time to a state of pluripotency by expressing appropriate transcription factors (the Nobel Prize in Medicine 2012). Cell reprogramming can be externally induced via the delivery of transcription factors, naturally or in vitro [5,6]. Such a reversion of a differentiated to a pluripotent state is the idea behind cancer stem cell (CSC). The CSC theory proposes that among all cancerous cells, a few act as stem cells that reproduce themselves and sustain cancer, much like normal stem cells that renew and sustain organs and tissues of body. The process of somatic reprogramming using Yamanaka factors, where many of them are oncogenes, offers a glimpse into how cancer stem cells may originate. In particular, neurological cancers such as primary glioblastomas [7] and retinoblastomas [8] are resulting from dedifferentiation of the glial and photoreceptor (retinal) cells, respectively. In particular, retinoblastoma tumor cells lose their *photoreceptorness* and become malignant plastic cells, i.e., CSC [8]. Xu et al. also have shown that the cell of origin for retinoblastoma is a committed cone precursor – an almost terminally differentiated photoreceptor – that has lost Rb gene, and not a pluripotent progenitor [9,10]. Here, we use the example of photoreceptor mosaic of avian retina to shed light on the following questions: **(Q1)** how cell intrinsic dynamics and microenvironmental factors coordinate during development to produce organized tissues such as photoreceptor mosaics [11]; and, **(Q2)** why the differentiated mosaic is so stable or how probable is the reversal of retinal tissue back to a pluripotent one, e.g., retinoblastoma.

The theory behind cell differentiation has been formalized in a metaphorical way by C. H. Waddington [12–14], which allows for developing a dynamical systems framework for modeling single-cell fate decisions [15–17]. Waddington has depicted the developmental process as a series of cell decisions that can be represented as bifurcations towards a differentiated state/phenotype. Practically, cell states, also called microstates, can be viewed as a vector of molecular expressions which are experimentally measured via high-throughput omics data, FACS, immunohistochemistry markers, etc. [14,18,19]. We note that such cellular microstates are technically different from the classical statistical mechanics definition of microstates. In general, differentiated states can be viewed as the fixed point(s) of microstate attractors [20,21]. Typically, these states can be associated with an appropriate probability distribution peaked around the fixed point, which is the deepest point of the valley in the Waddington potential. The situation is much more intricate in the case of pluripotent, stem cell-like states, where Waddington has also depicted them as attractors around a fixed point. However, this has been recently challenged by Furusawa and Kaneko [22], where they have shown that stem cell-like attractors can be viewed as limit-cycle attractors and not as stable fixed points. Biological observations of dynamic variability of single cells within pluripotent cell populations distinguish between pluripotency as a molecular state and pluripotency as a function, indicating that a pluripotent state is not unique but rather appears to be compatible with a wide variety of interchangeable molecular microstates (patterns of gene or protein expressions) [23,24]. In this view, pluripotent cells independently explore a variety of molecular expression states, and this phenotypic/state exploration transiently primes each individual cell to respond to a range of various differentiation-inducing stimuli, depending upon its instantaneous molecular state [7]. Finally, Waddington theory does not take into account cell sensing and the corresponding interactions that take place in a tissue. The aforementioned limitations of Waddington approach require further theoretical development to answer the questions **(Q1)** and **(Q2)**. In order to answer **(Q1)**, in this paper, we employ the recently proposed the Least microEnvironmental Uncertainty Principle (LEUP) – which is essentially a statistical mechanical theory for cell decision-making [25–27] – and apply it to the problem of cell differentiation. The LEUP is inspired by the theories of Bayesian brain hypothesis [28], the free-energy principle [29], and other dynamic Bayesian inference theories that try to explain human brain cognitive dynamics. Similar ideas have been also proposed in the influential work by W. Bialek [30]. Similar to these theories, the LEUP is based on the premise that cell internal molecular networks adapt to the sensed microenvironmental data and subsequently determine the relevant decisions. In turn, cells are encoding sensed information in genetic, epigenetic, translational, or transcriptional levels, where different timescales are related to each of these encoding levels, depending on the persistence of the microenvironmental stimuli. The reason to propose a theory such as the LEUP is the fact that the complexity of molecular networks does not allow us to know the exact involved dynamics, thus we use the LEUP as kind of dynamic Bayesian inference to circumvent this complexity and to make predictions. The LEUP theory also implies a decrease in the local microenvironmental entropy of the cell decision-maker, which biologically translates into the actions of differentiating cells which lead to more organized tissues during development.

The central cellular process related to the LEUP is cell sensing. Cells can acquire knowledge about their microenvironment by various sensing mechanisms such as binding of their receptors to diffusible ligands [31], pseudopodia extension [32], mechanosensing [33], proton-pump channels [34], gap junctions, etc. Cells can sense rapid changes of their milieus, where they mainly exploit two ways to decrease sensing errors: **(1)** by increasing the number of receptors or the responses of the downstream signaling pathways [35]; and/or, **(2)** by increasing sensing area [36]. The latter can span from the resting cell size to extensions via pseudopodia, blebbing, and other cell size regulation mechanisms. Pseudopodia or blebs can act as sensors, via a pressure sensing mechanism mediated by Piezo channels, allowing cell to decide when and where to migrate [37]. In this regard, we further specify our second question **(Q2)** as: can one calculate the limits of the cell sensing radius that ensure the robustness of differentiated tissue spatial order?

To answer **(Q2)**, we develop a thermodynamic-like theory using the tools of stochastic thermodynamics for a generic cell differentiation process. The main biological assumption is that differentiated cell can reverse to an undifferentiated state, for instance, by the process of carcinogenesis, see also Refs. [8,9,38]; a recent review on the latest advances in research on the process of dedifferentiation both at cell and tissue levels can be found in [39]. Stochastic thermodynamics allows us to understand the conditions that, even though single-cell dedifferentiation is possible (microscopic reversibility), the system (tissue) is still able to be robust and it maintains its spatial order and differentiation integrity (macroscopic irreversibility). Stochastic thermodynamics is a suitable tool for systems where small scale dynamics matters (e.g., soft matters, active matters, and biological systems), in such cases the higher-order moments dominate [40–44]; for different experimental applications of stochastic thermodynamics, see Refs. [45,46]. In addition, to formulate the laws of thermodynamics in the mesoscopic scale (specifically, at the level of trajectory) has also been investigated recently [47]. To describe cell-level dedifferentiation process in the context of stochastic thermodynamics, we apply the fluctuation theorem, specifically the Crooks’ relation [48,49]. The fluctuation theorem, which can be considered as the heart of stochastic thermodynamics, initially had been derived to explain how irreversibility at macroscopic level emerges from the underlying reversible dynamics and to estimate the probability of the violation of the second law of thermodynamics within a short amount of time for systems at small scales [50–52].

By combining elements of stochastic thermodynamics with the LEUP, we aspire to approach questions **(Q1)** and **(Q2)** within the context of avian photoreceptor mosaic. This paper is organized as follows: in Sec. 2, we review the basic concepts of the LEUP and its connection to statistical mechanics. In Sec. 3, we demonstrate how optimal microenvironmental sensing is associated with differentiated tissue spatial configuration, therein, we examine our theory using the data obtained from the (avian) photoreceptor mosaic. We apply the fluctuation theorem to cell differentiation in Sec. 4 and demonstrate the thermodynamic robustness of this process in the case of the avian retina development. In Sec. 5, we show how cell sensing radius and total entropy production are related and determine the limits of the sensing radius in order the tissue development to be robust. Finally, we conclude and discuss our results in Sec. 6.

## 2. Least microEnvironmental Uncertainty Principle

Cell collects information from its microenvironment and based on that it takes actions (phenotypic decisions). In other words, cell reacts to the environmental information by changing its own state. By adopting the notations of Ref. [25], we denote internal variables (such as gene expression, RNA molecules, metabolites, etc.) of the *n*-th cell as *x*_*n*_ and external variables (such as chemical signals, ligands, homotypic or heterotypic interactions, cellular densities, translational proteins, etc.) of the *n*-th cell as *y*_*n*_. The latter includes all the different extrinsic variables within the interaction radius of the *n*-th cell, i.e., *y*_*n*_ = {*y*(*r*) : *r* ∈ (*r*_*n*_, *r*_*n*_ + *R*]}, where *r*_*n*_ is position vector of the *n*-th cell, and *R* is the maximum microenvironmental sensing radius of the *n*-th cell. Cell sensing radius is a tunable variable controlled by cell and is regulated by various biophysical mechanisms, as stated in Introduction. This is precisely what *R* is modeling, where it has been considered as an intrinsic cell variable dictated by the LEUP dynamics. We also note that we have assumed a sensing radius can be identified with the typical interaction radius considered in agent-based models. In Fig. 1, we have shown the schematic diagram of the microenvironment of a differentiated cell as is being composed of both pluripotent progenitors and differentiated cells.

**Figure 1.**
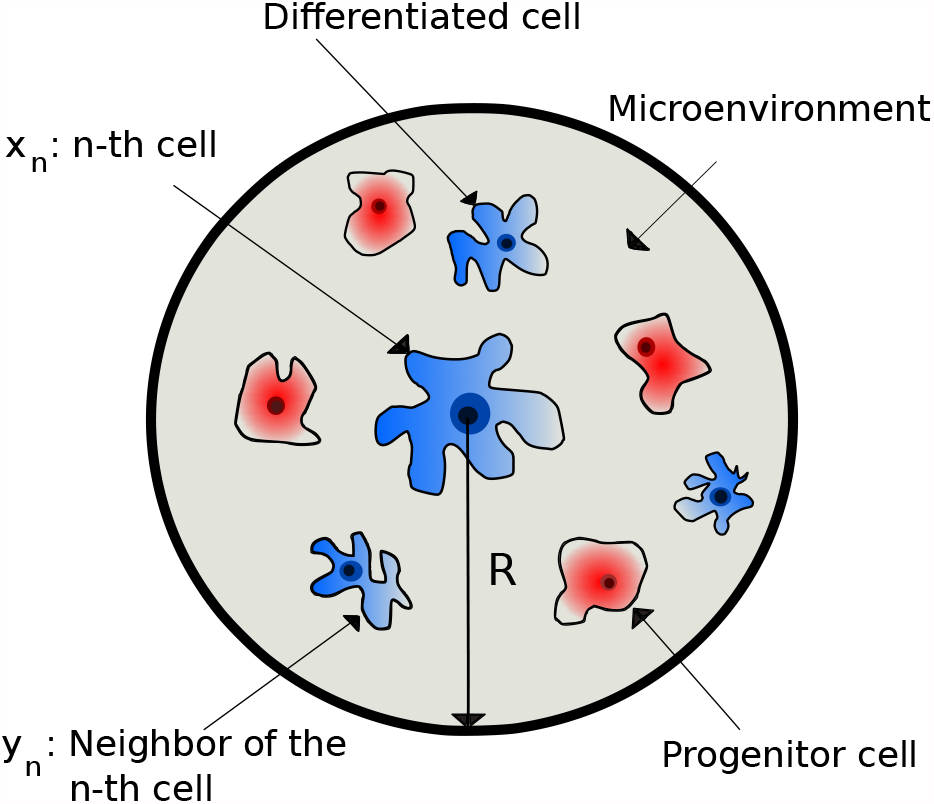
The schematic diagram of the microenvironment of a differentiated cell, where *R* is the maximum microenvironmental sensing radius. Microenvironment is being composed of a distribution of pluripotent progenitors and differentiated cells.

We consider behavior of cell, as a Bayesian decision-maker that reacts to its microenvironment [25,53], such that

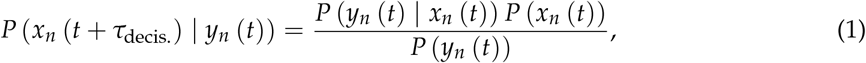

where *τ*_decis._ is the time needed that cell makes a decision. The distribution *P* (*y*_*n*_ (*t*) | *x*_*n*_ (*t*)) is the probability of microenvironmental information/data being collected by cell at time *t*. In other words, it is the probability that cell perceives all other cells, chemicals, and nutrients in its surrounding. The distribution *P* (*x*_*n*_ (*t*)) is the prior probability of the cell current internal states. If these two distributions are multiplied and be divided by the probability of external states *P* (*y*_*n*_ (*t*)), then (1) implies that the resulting quantity is the posterior probability distribution of internal states *P* (*x*_*n*_ (*t* + *τ*_decis._) | *y*_*n*_ (*t*)). The latter describes the most likely decision to be made by cell over the internal variables after processing the available information in the time period of *τ*_decis._.

As stated in Introduction, we assume that the cell prior is continually being updated by the previous time step posterior, in the sense of Bayesian learning. We also have assumed a *perfect* transfer of the prior to the next time step posterior, although this process is highly noisy. Now, by taking the logarithm of (1) and integrating over the joint probability distribution *P* (*x*_*n*_ (*t*), *y*_*n*_ (*t*)), for small decision times, we obtain:

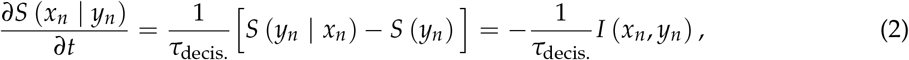

where for simplicity we have dropped *t* as an argument and *I* (*x*_*n*_, *y*_*n*_) is the mutual information. The above equation reaches equilibrium when the mutual information vanishes. Now, we make the first crucial assumption of our work **(A1)** that the cell decision time is much smaller than the asymmetric division time, i.e., *τ*_decis._ ≪ *τ*_div._. This can be justified as cell division which is a prerequisite for differentiation takes around ∼ 24*h*; on the other hand, the relaxation timescale of the molecular networks responsible for deciding a new state is much shorter (∼ 1*h*). This implies that the dynamics of internal variables *x*_*n*_’s are much faster than the microenvironmental dynamics of *y*_*n*_’s. According to this *timescale separation*, we can assume that variations of *S* (*y*_*n*_ | *x*_*n*_) and *S* (*y*_*n*_) belong to the slow manifold of the system. In turn, we observe that in order Eq. (2) reaches an equilibrium, the microenvironmental entropy sensed by cells *S* (*y*_*n*_ | *x*_*n*_) should be inevitably a decreasing quantity with respect to time. This fact can be interpreted as the entropy of cell sensing distribution should become more *focused* and more *independent* of the microenvironment as time goes on, since this information – microenvironmental data – has already been encoded in the cell prior. The latter is valid within biophysical context because of the following reasons: first, the collection of microenvironmental data, that is, the precise evaluation of *P* (*y*_*n*_ | *x*_*n*_) is energetically expensive, thus it is not favorable as cells would like to be energetically efficient; and second, fully differentiated cells are perfectly adapted to their microenvironment and the corresponding fluctuations implying an optimal prior distribution *P* (*x*_*n*_).

Using the timescale separation of **(A1)**, we formulate a variational problem concerning the maximization of entropy of cell microstates *S* (*x*_*n*_) in order to find an optimal prior *P* (*x*_*n*_), which is constrained by the current *local* microenvironmental entropy 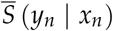 that corresponds to a random sample of size *R* of the fully differentiated tissue of interest. We note that due to **(A1)**, cell “solves” the entropy maximization by sensing an almost time-invariant microenvironmental entropy. We consider the definition of entropy in terms of probability distributions as *S* = −Σ_*i*_*P*_*i*_ ln *P*_*i*_ (Shannon entropy), where in the case of continuous set of states the summation symbol is to be replaced by an integral. Thus, our variational problem reads

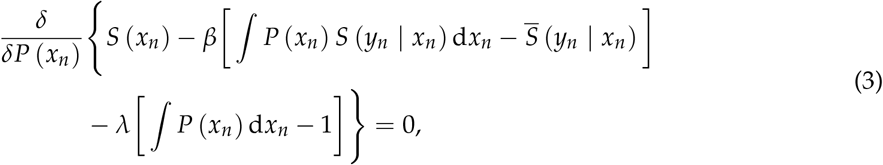

where *δ*/*δP* (*x*_*n*_) is the functional derivative. We have introduced two Lagrange multipliers, namely, *β* and *λ*. The parameter *β* is related to the current local microenvironmental entropy 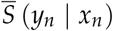 and *λ* is a constraint which preserves the normalization of probability. We note that other knowledge about the system, such as an explicit model in terms of statistical observables, can be incorporated as additional constraints. The solution of Eq. (3) then reads

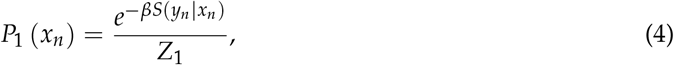

where 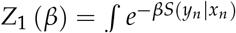d*x*_*n*_ is a normalization constant. We note that the parameter *β* quantifies the compliance of cell to the LEUP which is related to the ability of cells to sense their microenvironment and convert the perceived effects into biological signals. In the next section, we elaborate on the relation of *β* and microenvironmental sensing.

Now, we want to generalize (4) to a biologically relevant scenario of cell differentiation. It is well known that differentiation takes place during asymmetric divisions of pluripotent progenitor cells [54]. Let us assume that *µ* ∝ *τ*_div._ is the asymmetric division probability of a pluripotent cell. In a *mesoscopic* microenvironmental ensemble of *N* (*y*_*n*_ | *x*_*n*_) cells around the *n*-th cell, we want to calculate the probability of the central *n*-th cell to change its phenotype. This probability reads as

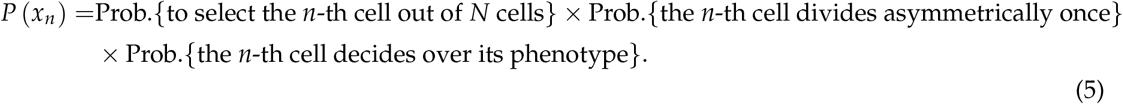

The latter probability happens according to the LEUP and it is the same as (4). Now, we assume that the probability of asymmetric divisions within an ensemble of *N* cells follows a *Poisson distribution* Pois[*µN* (*y*_*n*_ | *x*_*n*_)], which is a limit of the Binomial distribution B[*N* (*y*_*n*_ | *x*_*n*_), *µ*] for a small proliferation probability *µ*. Putting all these together, we can write:

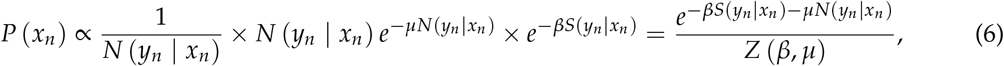

where the normalization factor of *P* (*x*_*n*_) is defined as 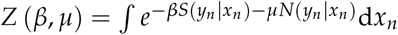.

We illustrate the above with a concrete example: if the microenvironmental probability distribution sensed by the *n*-th cell follows a Gaussian distribution with the variance 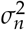, assuming that *y*_*n*_ is scalar, and hence 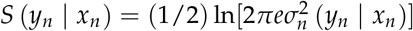, and by considering *µ* = 0, then (6) reduces to

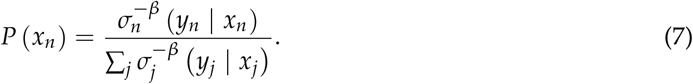

Now, we are in a position to establish an explicit formula for cell internal entropy. To this end, by exploiting (6), we obtain

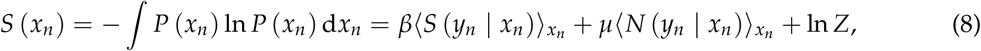

where ⟨ … ⟩ denotes the expectation value. By making usage of the notions which are developed within the context of thermodynamics, we can also define phenotypic internal energy sensed by cell as

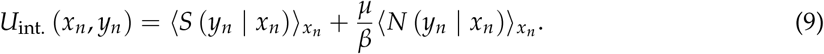

We note that for *µ* = 0, the internal energy is the same as entropy for a particular realization. Thus, (6) resembles the Boltzmann distribution.

Interestingly, since the microenvironmental entropy *S* (*y*_*n*_ | *x*_*n*_) acts as an effective internal energy, it is expected to be a decreasing quantity in time. This result is in agreement with the discussion which follows (2). Thus, the LEUP is consistent with its premises.

### 2.1. The LEUP and known statistical results

The steady-state distribution of (7) which is derived by the LEUP, under the assumption of Gaussian distributed microenvironment and *µ* = 0, can be associated with different established statistical results. Here, we distinguish several cases for specific values of *β*, which have particular meanings in statistics. For instance, when *β* = 1, we reproduce distributions proportional to the well-known Jeffreys prior, see also Ref. [25]. Jeffreys prior is known as the most typical uninformative prior used in Bayesian inference [55].

Interestingly, if we assume that *β* = 2, then we recover the so-called *minimum variance estimator* when fusing multiple scalar estimates. Following [56], let us assume extrinsic variables *y*_1_, *y*_2_, …, *y*_*n*_, such as ligand concentrations, cell densities, etc., that are pairwise statistically uncorrelated and are normally distributed, i.e., 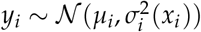, where 1 ≤ *i* ≤ *n*. In turn, we assume intrinsic variables *x*_1_, *x*_2_, …, *x*_*n*_, that correspond to cellular sensors and downstream process of the aforementioned microenvironmental variables. We define the average sensed microenvironment as the average of the extrinsic signals *y*_*i*_’s weighted by the distribution of the cell sensors *x*_*i*_’s, i.e., 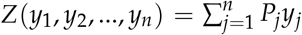, where 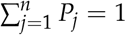. The distribution of internal variables *P*_*i*_ that minimizes the variance, aka noise of the sensed microenvironment, is given by the following formula:

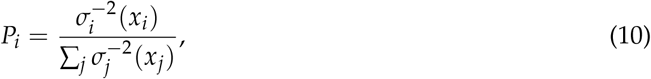

which has the same form as the steady state of the LEUP (7), for *β* = 2. In the following, we connect the latter result with a specific cell sensing scenario.

## 3. Minimal cell sensing noise is associated with differentiated tissue spatial configuration

In this section, first we show how the LEUP is connected to cell sensing with respect to receptor-ligand binding. Then, we apply our theory to avian photoreceptor mosaic and fit the parameter *β* to recover the photoreceptor percentages in retina.

### 3.1. How does the LEUP relate to cell sensing?

In this subsection, we show the relevance of the LEUP within the context of cell sensing mechanisms. Here, we focus on the receptor-ligand sensing apparatus of cell, where complexes are formed at cell membrane and subsequently internalized via endocytosis [31]. This sort of sensing mechanism is also relevant in the case of photoreceptors [57]. Receptors act as sensors for specific microenvironmental molecules (ligands) that bind together with a certain affinity to form complexes. Often different ligands may bind to the same receptor, for instance, the Notch-Delta-Jagged system where Notch receptor can bind to either Delta or Jagged molecules [58,59]. The sensed information can be quantified by the concentration of internalized complex molecules. For simplicity, we denote *x* as receptor concentration at the cell membrane and *y*_1_ and *y*_2_ as the corresponding ligands concentrations, which we assume that they are statistically independent. Moreover, the ligands concentrations are consumed by cells, since they bind to receptors and therefore depend on the concentration *x* (we omit this dependence for the sake of simplicity). Typical dynamics of such systems reads as

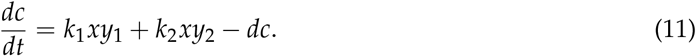

Under the assumption of fast decay rates *d* ≫ 1, which is consistent with **(A1)**, the system behaves as in a steady state, that is,

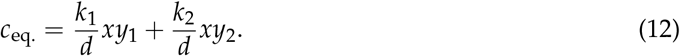

The terms, (*k*_*i*_/*d*)*x, i* = 1, 2, define the percentage of receptors bound to ligands *y*_*i*_’s. In the context of the LEUP, (*k*_*i*_/*d*)*x* corresponds to *P*(*x* | *y*_*i*_), which under the assumption of Gaussian microenvironment – in a steady state – is:

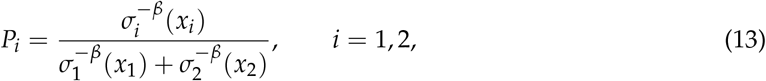

which for *β* = 2 coincides with (10). Taking all the above together, we translate the receptor-ligand cell sensing system as a linear combination of complex formation estimates *y*_*i*_’s, where the LEUP probabilities *P*_*i*_’s are the corresponding coefficients/proportions of complexes bound to ligands *y*_*i*_’s, that is,

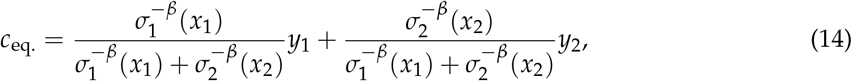

which coincides with the definition of *Z*(*y*_1_, *y*_2_, …, *y*_*n*_), for *n* = 2, defined in the previous section.

At this point, we want to do a sanity test for our results. For simplicity, we consider a single type of ligand concentration *y* that binds to a receptor of concentration *x*. Then, the complex formation dynamics is described as

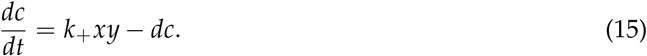

At the equilibrium, the steady-state complex concentration *c*_eq._ reads

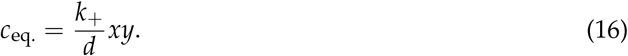

As before, we define the probability *P*_*x*_ = (*k*_+_/*d*)*x* as the proportion of binding receptors. Then from (16), we observe that:

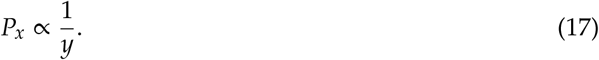

The ligand concentration, *Y* = ⟨*ϒ*⟩, is the expected value of the number of ligand molecules within the cell volume, which is denoted as *ϒ*. Using arguments similar to that of Berg and Purcell in their seminal paper [35], the ligand molecules are diffusing and therefore we can assume *ϒ* follows a Poisson distribution, thus 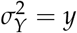, where the ligand variance is denoted as 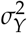. Now, by combining the latter with (17), we recover our non-normalized the LEUP result for *β* = 2:

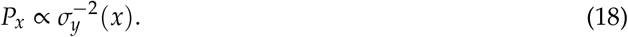

This specific value of *β* complies with the *perfect monitoring/sensing* assumptions (for details, see [35]). We note that in (18), we have explicitly denoted the dependence of diffusible ligand concentration on the receptor concentration *x*. As a side remark, if we change the receptor-ligand binding term by introducing, for instance, finite number of receptors or covalent bonds, then the parameter *β* will be modified.

The above result is pivotal since it connects the cell fate decision-making with optimal microenvironmental sensing in terms of minimization of sensing noise, which results into a specific spatial phenotypic distribution. In the following, we explore the validity of these results in the case of the avian photoreceptor mosaic.

### 3.2. Application: predicting the cone color distribution in the avian photoreceptor mosaic

In this subsection, we focus on a particular multicellular system, namely, the photoreceptor cone mosaic of the avian retina. Retina plays a vital role in the visual perception of vertebrates allowing for light sampling and signal transfer to the corresponding optic nerves. The photoreceptor layer is responsible for the efficient light collection and color encoding of the surrounding structures. The avian retina possesses one of the most sophisticated cone photoreceptor systems among vertebrates [60]. Birds have five types of cones including four single cones, namely, green, red, blue, and violet, and a double cone, whereas in humans there are only three: red, green, and blue. In the case of birds, these cones are ordered on the retina in a non-perfect hexagonal manner. The five avian cone types exist as five independent spatial mosaics that are all embedded within a single monolayered epithelium along with a population of rod photoreceptors. In Ref. [60], the spatial organization of these photoreceptor mosaics has been analyzed, where the authors have found that the five cone types are present in the characteristic ratios at the latest stage of development, as is illustrated in Fig. 2.

**Figure 2.**
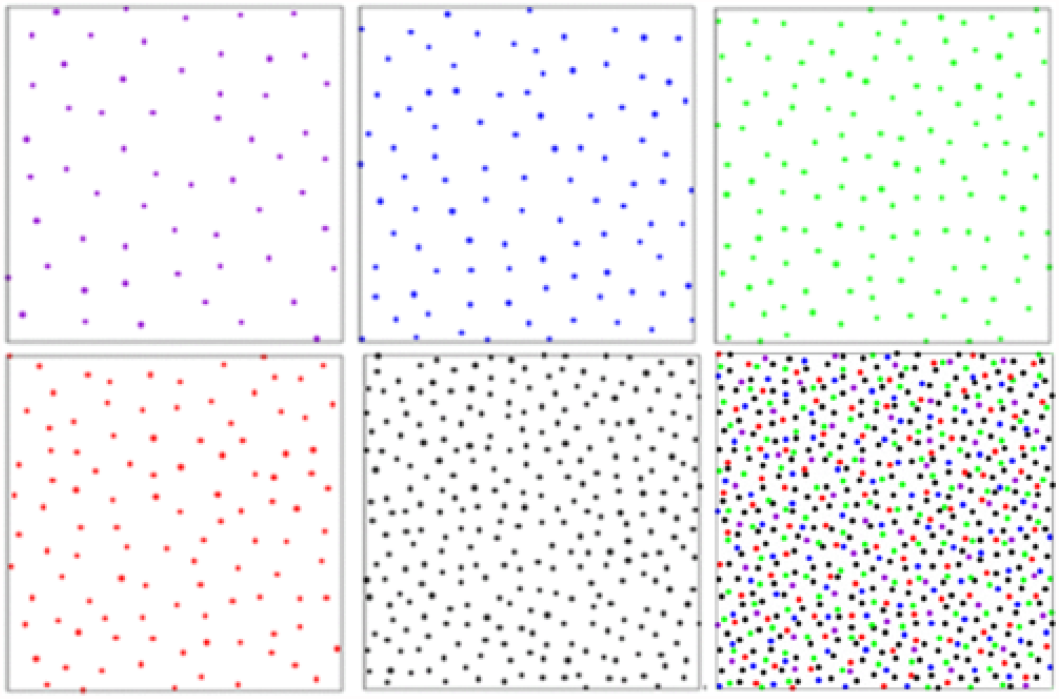
The experimentally obtained chicken cone photoreceptors arrangement in space. In the upper panels, the configurations from left to right correspond to violet, blue, and green species, respectively. In the lower panels, from left to right, the configurations correspond to red, double species, and the overall pattern, respectively. The figure is adapted from [61].

Our goal is to reproduce the ex vivo observed cone distributions of [60] by using the aforementioned LEUP-driven results. In particular, we intend to fit the parameter *β* in formula (7) using the photoreceptors data. Therefore, let us consider we have *n* = 5 different cone states *x*_*i*_’s that correspond to the expressions of different opsins, *i* = *g, r, b, v, δ* (*g* stands for green, *r* for red, *b* for blue, *v* for violet, and *δ* for double). Moreover, we define the corresponding cone occurrence probabilities *P*_*i*_ = *P*(*x*_*i*_) as cone photoreceptors percentages in retina. In turn, we denote *σ*_*i*_ as the standard deviation for a given spatially local neighborhood related to the Nearest Neighbor Distribution (NND). In Ref. [60], the authors have exploited the Delaunay triangulation in order to calculate the NND for each cone and they have reported the first and second moments of it. In order to compare the LEUP color distributions against the ex vivo ones, we employ the Kullback-Leibler divergence. In turn, we numerically search for the *β*’s which appropriately fit the physiologically observed photoreceptor fate ratios. In the left panel of Fig. 3, we have depicted the Kullback-Leibler divergence, i.e., *D*_*KL*_ = ∑_*i*_ *P*_*i*_ ln(*P*_*i*_/*Q*_*i*_), as a function of *β*, where *i* denotes different cone cells, and *P*_*i*_ and *Q*_*i*_ correspond to the probabilities obtained from the experimental data and the LEUP, respectively. The plot demonstrates that *D*_*KL*_ has a minimum of ≈ 0.004 at *β* ≈ 1.754. We have compared the ex vivo observed cone distributions to the LEUP, in the right panel of Fig. 3. Interestingly, the *β* values that provide the best fit are in the range 1.378 ≤ *β* ≤ 2.136, where the values of the Kullback-Leibler divergence are in the order of magnitude 10−3. Thus, it demonstrates that the values of *β* ≈ 2 explain the observed data, implying that the avian cone mosaic distribution is associated with an optimal cell sensing process that minimizes the corresponding noise.

**Figure 3.**
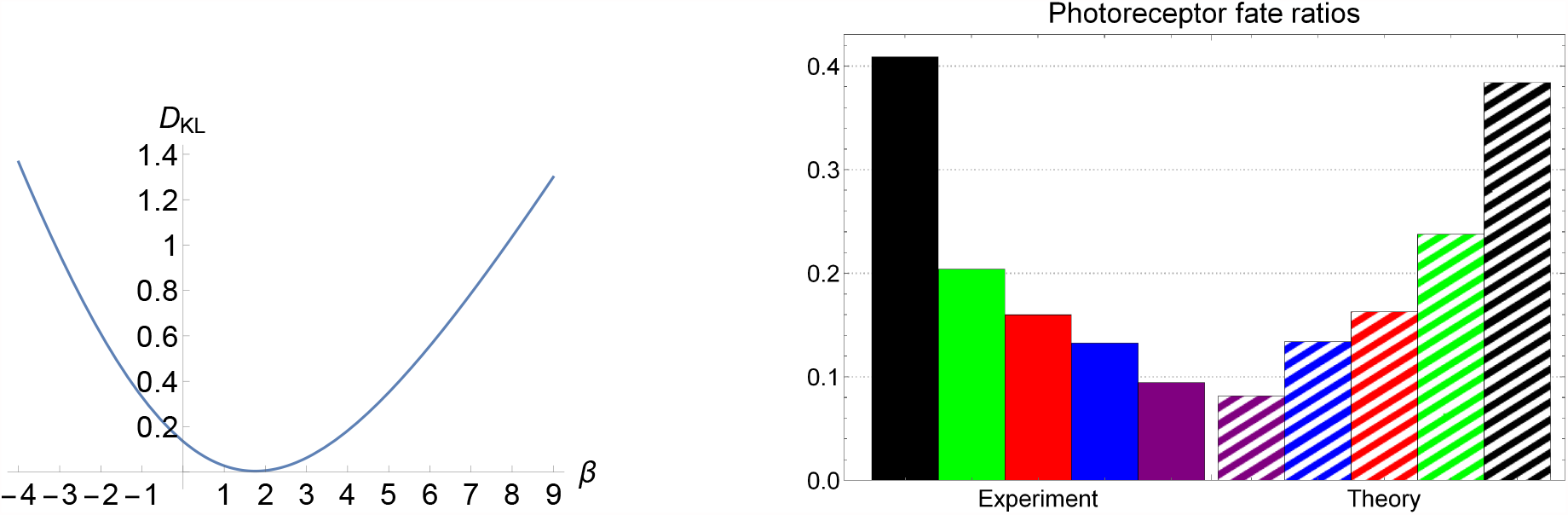
In the left panel, the Kullback-Leibler divergence *D*_*KL*_ is depicted as a function of *β. D*_*KL*_ reaches its minimum of ≈ 0.004 at *β* ≈ 1.754. The experimentally obtained photoreceptor fate ratios is compared to the LEUP for this particular *β*, in the right panel.

## 4. Thermodynamic robustness of differentiated tissues: the fluctuation theorem

In this section, we determine the thermodynamic constraints of two coarse-grained cell states that correspond to pluripotent (*s*) and differentiated (*d*) states. Then, we apply the obtained results to the particular case of the avian cone cells differentiation and the formation of photoreceptor mosaics. To this end, first we show how microstates (internal variables) are related to the microenvironmental information and heat transfer (cell metabolism).

In this context, a cellular *microstate* corresponds to a cell phenotype that lives in a tissue, which could be gene expression, RNA molecules, receptor distribution, etc. In other words, microstate gives information about the internal states of cell. We label these internal variables as *x*_*s*_ and *x*_*d*_ corresponding to pluripotent and differentiated cells, respectively. We define a cellular *macrostate* as a statistical observable (e.g., average) of a cell microenvironment that involves multiple cells of different phenotypes. Macrostates contain information about external variables which are labeled as *y*_*s*_ and *y*_*d*_. The macrostate *y*_*s*_ is assumed to describe a microenvironment of pluripotent progenitor or stem cells characterized by the microstate *x*_*s*_; the macrostate *y*_*d*_ accordingly describes a microenvironment of differentiated cells characterized by the microstate *x*_*d*_. We denote the number of pluripotent cells neighboring to a cell with microstate *x*_*s*_ as *N*(*y*_*s*_ | *x*_*s*_) and the number of differentiated cells neighboring to a cell with microstate *x*_*d*_ as *N*(*y*_*d*_ | *x*_*d*_). The total number of pluripotent and differentiated cells inside system are denoted as *N*(*s*) and *N*(*d*), respectively.

Now, based on (6), we can write the probability of cell to be in the microstate *x*_*s*_ with the corresponding macrostate *s* as

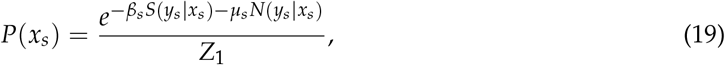

and the probability of being in the microstate *x*_*d*_ with the corresponding macrostate *d* as

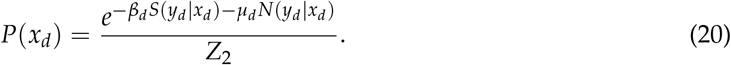

It is known that for a system which is coupled to a set of heat baths and is in a time-symmetrically driven nonequilibrium state, the Crooks’ theorem is applicable [48,49]. In this case, the dynamics follows Brownian motion, i.e., there are no extreme *jumps* in the system state. For any trajectory to be initially at *x*_*s*_(0) and is going through microstates *x*_*s*_(*t*) over time *τ*, the Crooks’ theorem implies that [49],

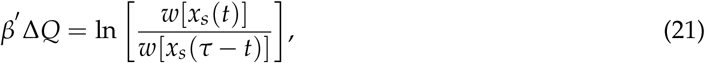

where *β ′* ≡ 1/*T*, which *T* is the temperature of heat bath, Δ*Q* is the total heat released into bath over the course of *x*_*s*_(*t*), and *w*[*x*_*s*_(*t*)] is the probability of trajectory *x*_*s*_(*t*). Eq. (21) demonstrates that when there is a forward state change, system loses heat to reservoir and in the case of a time-reversed path, there is a heat gain from reservoir; this, in turn, implies that a forward trajectory is more probable than a time-reversed one, thus, Eq. (21) substantiates a relation between heat and irreversibility at a microscopic (cell) level. This conclusion is in agreement with the ideas presented in [7]. The authors of the mentioned paper have considered differentiation as a series of reversible transitions through many microstates (we shall use this notion when discussing differentiation at a tissue scale in the following), where stem cells exhibit reversible oscillations until an attractor drives them towards a differentiated state. Within this picture, dedifferentiation is more likely to occur only on a small scale with a low probability, as a series of microstate transitions. We also note that, as is demonstrated in Ref. [49], Eq. (21) is also valid for other kinds of steady-state probability distributions besides the classical Boltzmann distribution. In our case, such Brownian jumps are interpreted as changes in phenotypes and the associated heat losses are assumed to be due to potential changes in cell metabolism. Another important heat loss contribution comes from the cell divisions, which are required in the differentiation process [62]. However, there are other minor heat loss sources that are disregarded since they act on shorter time scales, such as physical friction, changes in the cytoskeleton, etc.

Our goal here is to establish a microreversibility relation for a general differentiation process (the details of derivation can be found in Appendix). To this end, first we fix the starting point of a differentiation trajectory as *x*_*s*_, which is a microstate realization that belongs to a pluripotent cell attractor/set, and the ending point as *x*_*d*_, that is a member of the differentiated phenotypes set (see Fig. 8 in Appendix). The Crooks’ theorem in (21) is valid for a single differentiation trajectory that connects a single pluripotent state to a particular differentiation state realization out of many. Therefore, first we average over all possible paths that connect a pluripotent and a differentiation state, and then we average over combinations of dedifferentiation paths that connect any pluripotent to any differentiation states, resulting into:

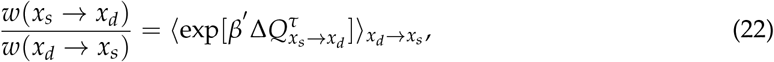

where *w*(*x*_*s*_ → *x*_*d*_) is the transition probability that system is found to be in the microstate *x*_*d*_ at time *τ*, given that system was initially in the microstate *x*_*s*_. The averaged version of the Crooks’ theorem in (22) has an important implication since it demonstrates that paths leading to differentiation and also dedifferentiation are many more than one.

Now that we have microscopic relation of (22) at our disposal, we can study macroscopic (tissue) consequences of that. At this point, we make the second vital assumption of our study **(A2)** that the tissue dynamics follows a Markov process. The probabilistic description of macroscopic states of two distinct cell types, from which one can understand the phenomenon of irreversibility at a macroscopic level, can be constructed as [49],

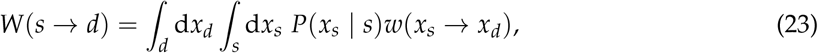

and

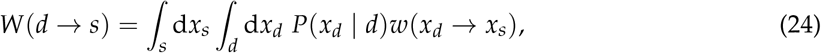

where *P*(*x*_*s*_ | *s*) is the probability that system to be in the microstate *x*_*s*_, given that it is observed in the macrostate *s*, and *w*(*x*_*s*_ → *x*_*d*_) is defined as before, where we have omitted *τ* for the notational convenience. The transition probability *W*(*s* → *d*) in (23) implies the likelihood that cell to be observed in the macrostate *d* while it was initially prepared in the macrostate *s*; accordingly, (24) is understood in the same fashion, i.e., the likelihood that a microenvironment being prepared in the state *d* to satisfy the microenvironment *s* after another time interval *τ*. These processes are illustrated in Fig. 4.

**Figure 4.**
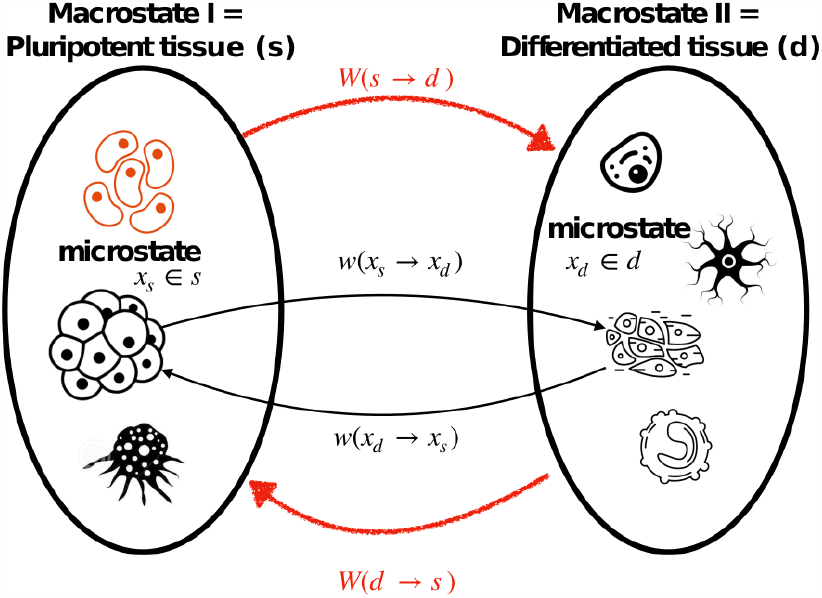
Microscopic/Macroscopic transitions between two distinct cell/tissue types.

By taking the ratio of (23) and (24), we obtain

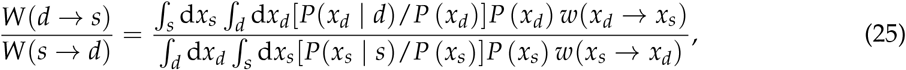

where we have multiplied and divided the numerator and the denominator by *P* (*x*_*d*_) and *P* (*x*_*s*_), respectively. We define the pointwise mutual information for individual trajectories as *i*_1_ = ln[*P*(*x*_*s*_ | *s*)/*P*(*x*_*s*_)] and *i*_2_ = ln[*P*(*x*_*d*_ | *d*)/*P*(*x*_*d*_)], and then by taking these definitions into account and replacing *P*(*x*_*d*_) by its corresponding relation in (20), (25) reduces to

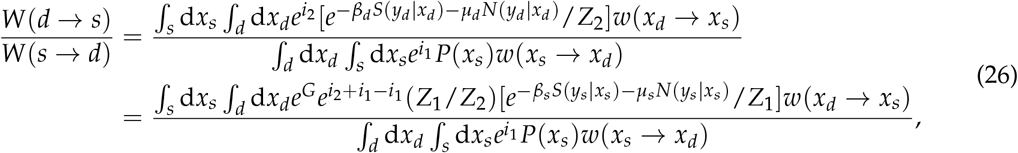

where

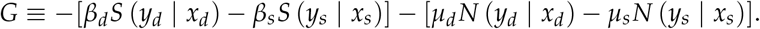

By exploiting (19) and (22), we can rewrite (26) as

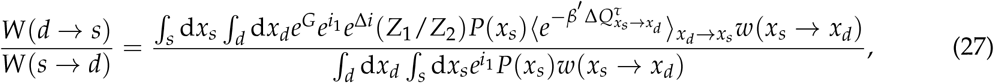

where Δ*i* ≡ *i*_2_ − *i*_1_

Now, (27) can be expressed in terms of the average over all trajectories from the ensemble of microstates *x*_*s*_ which correspond to the macrostate *s* to the ensemble of microstates *x*_*d*_ which correspond to the macrostate *d* while each path is weighted by its probability as

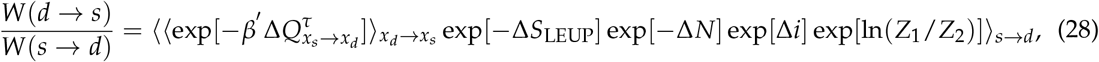

where

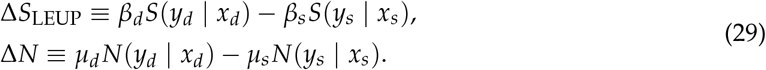

By rearranging (28) as

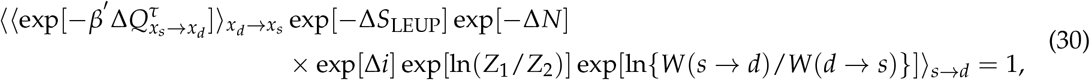

and using the Jensen’s inequality, i.e., exp[⟨*X*⟩] ≤ ⟨exp[*X*] ⟩, and the fact that *e*^*x*^ ≥ 1 + *x*, we arrive at

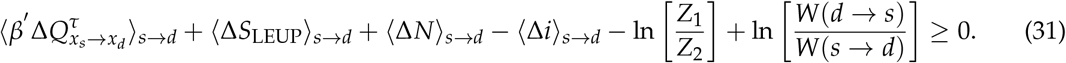

We note that if *s* and *d* correspond to the same identical classes, i.e., *W*(*d* → *s*) = *W*(*s* → *d*), then the last term of (31) vanishes. Now, by defining the remaining terms as *total entropy production*, that is,

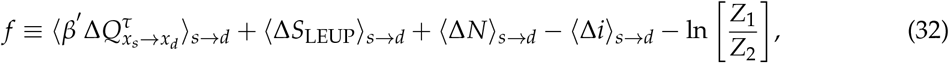

then (31) implies that total entropy production is always non-negative. In other words, (31) can be considered as a generalization of the second law of thermodynamics (see also Ref. [49]).

The above inequality is reformulation of the fluctuation theorem for tissue differentiation. It relates thermodynamic properties of system to the LEUP; and as is obtained under general assumptions, it is applicable to a general cell and tissue differentiation process. In the next subsection, we show how (31) implies the thermodynamic robustness of cell differentiation, in the particular case of the avian cone photoreceptor mosaics formation.

### 4.1. Application: differentiated photoreceptor mosaics are thermodynamically robust

In this subsection, we use the fluctuation theorem, culminated in (31), to demonstrate the robustness of the avian photoreceptor mosaics. By defining *s* → *d* as a pluripotent tissue differentiates to a more specialized one and denoting its corresponding forward transition probability as *p* _*f*_, i.e., *W*(*s* → *d*) = *p* _*f*_, and *d* → *s* as differentiated tissue dedifferentiates to the pluripotent one with its backward transition probability of *p*_*b*_, i.e., *W*(*d* → *s*) = *p*_*b*_, we can rewrite (31) in terms of *p* _*f*_ and *p*_*b*_ as

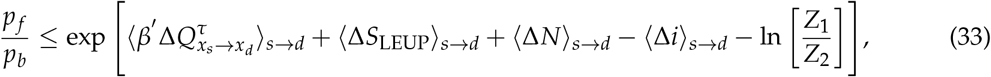

where we have already defined the exponent of the exponential in the right-hand side of (33) as total entropy production, see (32).

In order to obtain a thermodynamic constraint which ensures the robustness of differentiated tissue, first we assume that there exists a maximum forward transition probability from *s* to *d* in such a way that

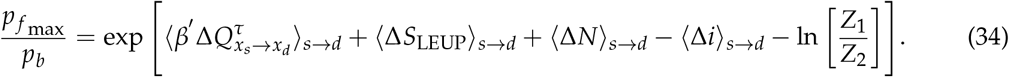

To simplify (34) more, we note that based on (8), we can write

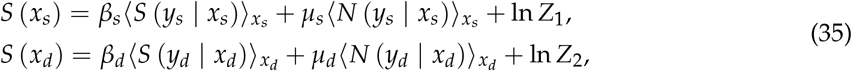

and then by subtracting these two and taking the average over all trajectories from *s* to *d*, we obtain

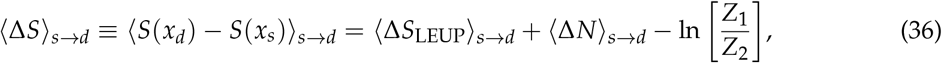

where we have used the definitions of Δ*S*_LEUP_ and Δ*N* as given in (29). Now, (34) can be written as

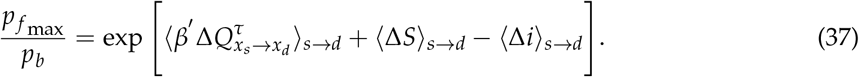

The immediate implication of (37) is that, differentiated tissue is thermodynamically robust if

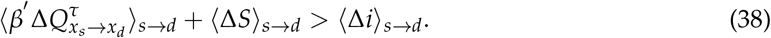

In the following, we present the avian cone cells differentiation and the corresponding photoreceptor mosaics formation as an example of which the above inequality is satisfied. In other words, we demonstrate the robustness of the avian retina development and the irreversibility of the time arrow in this particular process.

First, we note that as heat dissipation (the first term in the left-hand side of (38)) depends on metabolic pathways, this implies the crucial role of cell metabolism in the process of cell differentiation. In addition, consumption of glucose depends upon cell types. Progenitor or pluripotent stem cells use glucose as the primary metabolites for anaerobic glycolysis (fermentation) pathway: Glucose + 2 ADP + 2 Phosphate → 2 Lactate + 2 H^+^ + 2 ATP, where as a result of the breakdown of glucose to lactic acid the amount of energy around 109.4 kJ/mol is released [63]. Differentiated cells use glucose to produce carbon dioxide and water by using aerobic (respiration) reaction: Glucose + 6 O_2_ + 36 ADP + 36 Phosphate → 6 CO_2_ + 6 H_2_O + 36 ATP, where the released energy is around 2820 kJ/mol [63]. Thus, 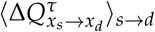 which is the total heat released during the journey from *s* to *d* is 2929.4 kJ/mol. As these reactions have taken place in *T* = 310 K, we have: 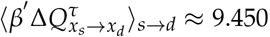, where we have dropped its dimension as we have set the Boltzmann constant to be 1, in this paper.

Now, in order to investigate (38) for the avian cone cells differentiation, we need to calculate the value of ⟨Δ*S*⟩ _*s*→*d*_ for this particular process. Within the context of photoreceptor cells, differentiated states are given by an array of five opsin expressions as *x* = (*x*_*g*_, *x*_*r*_, *x*_*b*_, *x*_*v*_, *x*_*δ*_) and the corresponding probability distribution of *P*(*x*) = (*P*_*g*_, *P*_*r*_, *P*_*b*_, *P*_*v*_, *P*_*δ*_). At this point, we assume – **(A3)** – that pluripotent state corresponds to a state which is compatible to an equiprobable distribution of opsins, that is, *P*(*x*_*s*_) = (1/5, 1/5, 1/5, 1/5, 1/5). As is discussed in Introduction, pluripotent states are not considered as attractive fixed point(s) but rather as oscillating attractors that *explore* the cellular state space [22]. Differentiated states *x*_*d*_’s are all the states that their *P*(*x*_*d*_)’s are equal to unimodal distributions centered at certain colors. Kram et al. [60] have reported different color percentages of the green, red, blue, violet, and double cone cells inside the avian retina (see Supporting Information of the mentioned reference). By assuming these numbers as the occurrence probabilities of the corresponding cones, we can write: *P*_*g*_ ≈ 0.204, *P*_*r*_ ≈ 0.160, *P*_*b*_ ≈ 0.133, *P*_*v*_ ≈ 0.094, and *P*_*δ*_ ≈ 0.409. Thus, we are able to calculate entropies of the individual cone cells and as a result that of *S*(*x*_*d*_) as

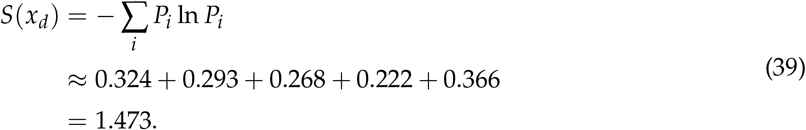

Now, the entropy difference between differentiated and stem cells reads

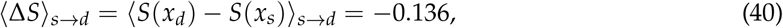

where *S*(*x*_*s*_) behaves as the entropy of a uniform distribution, i.e., 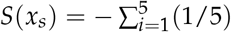; this is due to the fact that for pluripotent stem cells there are no yet color preferences.

In order differentiated tissue maintains its spatial order and integrity, (38) imposes a lower bound on heat dissipation (cell metabolism) as

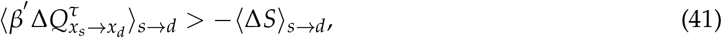

where we have set ⟨Δ*i*⟩ _*s*→*d*_ → 0 for simplicity. Inequality (41) is strongly hold for the values obtained here, that is, 9.450 ≫ 0.136. This implies that the development of the avian retina is highly robust. In other words, the arrow of time is almost irreversible in this process.

## 5. Cell sensing radius limits for robust tissue development

In this section, we derive a relationship between total entropy production and cell sensing radius in the particular case of progenitor cells differentiation into the avian cone photoreceptors. We calculate the limits of the sensing radius and, in the parametric space of the LEUP parameters, we suggest the biologically and physically acceptable regions for robust tissue differentiation.

In (32), we have introduced total entropy production as

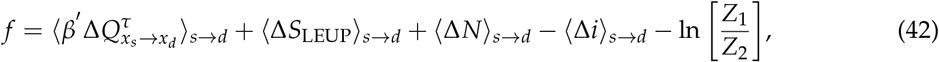

where thermodynamic properties of system are related to the LEUP quantities. In order to calculate ⟨Δ*S*_LEUP_ ⟩_*s*→*d*_, by assuming the microenvironmental probability distributions as Gaussians, we have

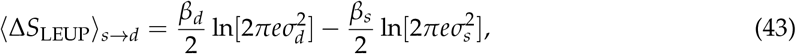

where we have used (29). The above can be simplified to

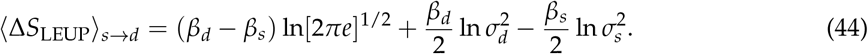

Now, we need to postulate the forms of 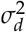 and 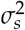 and their scaling with the corresponding cell sensing radius. Jiao et al. in [61] have found that the differentiated retina mosaic is hyperuniform, that is, 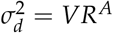, where *A* < *D*, and *D* = 2, 3; on the other hand, for progenitor cells, we assume – **(A4)** – a Poisson distribution, i.e., 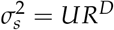. Here, *R* is the sensing radius, and *V* and *U* are the densities of the cone and progenitor cells neighbors, respectively. Plugging these formulas in (44) leads to

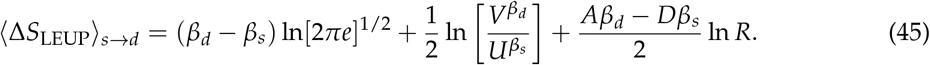

Another term of (42) which needs to be dealt with is ⟨Δ*N* ⟩_*s*→*d*_. From (29), we have

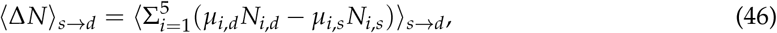

where *i* counts different types of cones: green, red, blue, violet, and double. We note that the population at each tissue reads as *N*_*i,j*_ = *ρ*_*i,j*_*R*^*D*^, where *ρ*_*i,j*_ is the density of microenvironment and *j* ∈ {*s, d*}. Thus, (46) reduces to

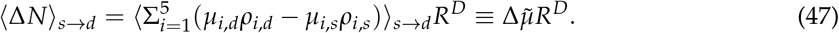

Putting all terms together, total entropy production (42) becomes

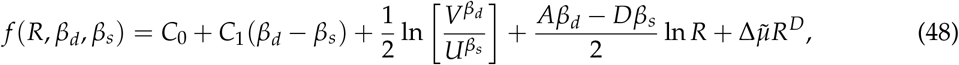

where we have assumed 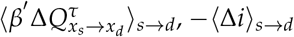 and −In (Z_1_/Z_2_) as constants and have grouped them together as *C*_0_, and *C*_1_ ≡ (1/2) ln(2*πe*).

Now that we have an explicit formula for total entropy production at our disposal, we can obtain the optimal sensing radius for which total entropy production reaches its extremum. To this end, we note that the first derivative of (48) vanishes at

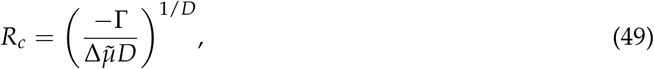

where Γ ≡ (*Aβ*_*d*_ − *Dβ*_*s*_)/2. When *R*_*c*_ is a positive quantity, then the inequality 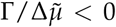 should always be satisfied. In order to determine the conditions for which *R*_*c*_ minimizes or maximizes total entropy production, we calculate the second derivative of (48) at (49), and obtain

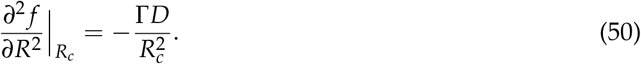

Thus, *R*_*c*_ minimizes total entropy production if Γ < 0 and it maximizes if Γ > 0.

In the left panel of Fig. 5, we have illustrated total entropy production (48) as a function of *R* for a particular set of the LEUP parameters which minimizes total entropy production based on (50). The values of the curve above the *R*-axis (gray line) are biologically and physically relevant as total entropy production is positive, i.e., it ensures the robustness of differentiation process. Moreover, this plot shows that as we move away from *R*_*c*_, (positive) total entropy production is rapidly increasing. The right panel of the figure illustrates *f* as a function of *R* and *β*_*d*_, where the acceptable regions lie above the gray plane. Interestingly, minimization of total entropy production occurs only for 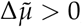, which assumes a decrease in the proliferative activity of differentiated phenotypes.

**Figure 5.**
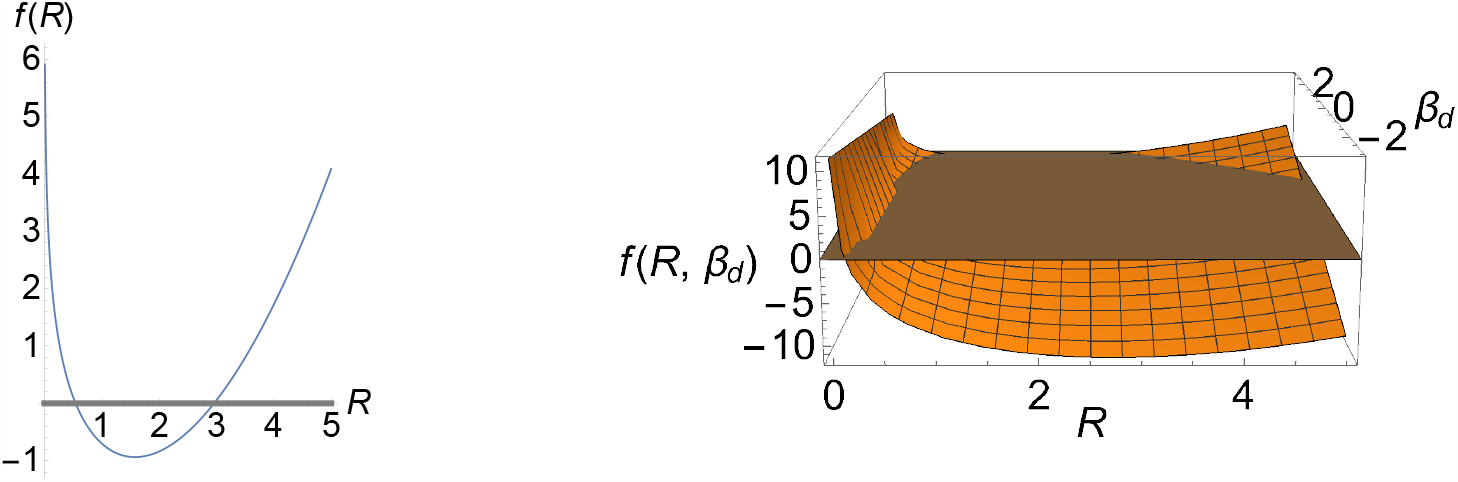
The left panel shows total entropy production as a function of sensing radius for *C*_0_ = −1, *β*_*d*_ = *β*_*s*_ = 3, *V* = *U* = 1, *A* = 1, *D* = 2, and 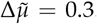. Based on (50), this specific set of parameters leads to a total entropy production which has a minimum of ≈ −0.937 at *R*_*c*_ ≈ 1.581. In the right panel, *β*_*d*_ is also treated as a variable. Due to the fact that total entropy production is always positive, the curve/surface above the gray line/plane is only biologically and physically acceptable.

In Fig. 6, the case for which total entropy production reaches its maximum is shown. This figure illustrates that (positive) total entropy production is bounded in this case. The immediate implication of this is tissue dedifferentiation is possible (negative total entropy production) for a large sensing radius. If we want to have a positive total entropy production, in order to avoid reversibility of differentiated tissue, we must have some sort of fine-tuning in order to restrict the values of *R* in such a way that they lead to a positive *f*. According to Bialek’s postulated biophysical principles [30], fine-tuning in Nature is not favorable. In the case of the avian photoreceptors, we can further constrain our parameters and identify further arguments to exclude this case.

**Figure 6.**
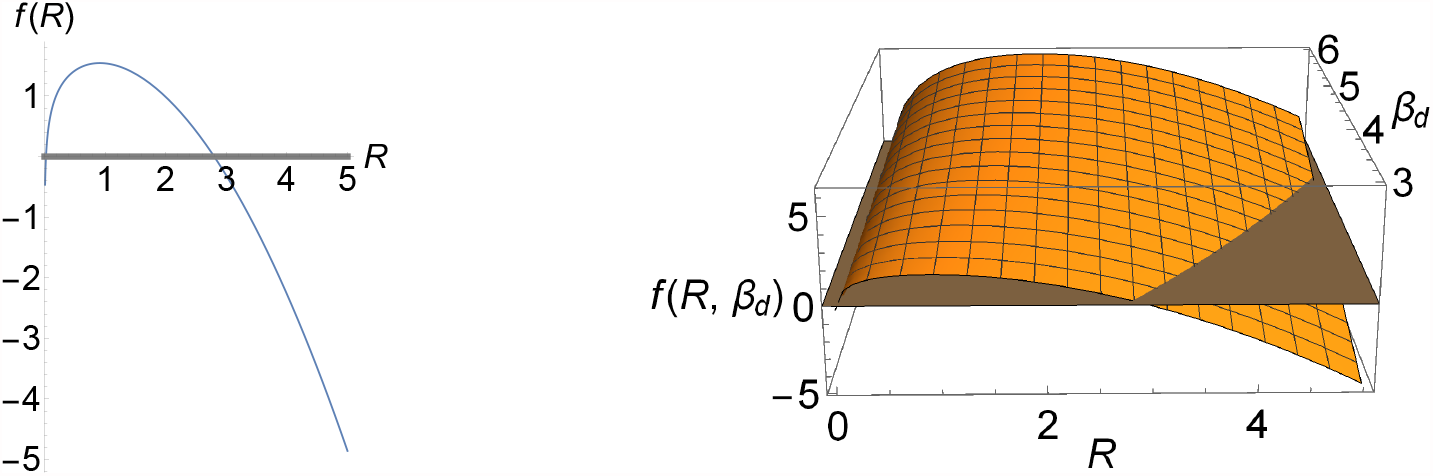
Total entropy production *f* as a function of *R* and (*R, β*_*d*_) is shown in the left and right panels, respectively. In the left panel, we have fixed the parameters as: *C*_0_ = −1, *β*_*d*_ = 3, *β*_*s*_ = 1, *V* = *U* = 1, *A* = 1, *D* = 2, and Δ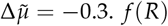 reaches its maximum of ≈ 1.542 at *R*_*c*_ ≈ 0.913. In the right panel, *β*_*d*_ is also considered as a variable. In both panels, the only biologically and physically acceptable regions lie above the gray line and the gray plane, as total entropy production should be positive; this makes total entropy production a bounded function in contrast to the case depicted in Fig. 5.

### 5.1. Application: the average sensing radius of the avian cone cell

In this subsection, we calculate the average sensing radius for each photoreceptor and by exploiting (48) we look for the regions, in the parametric space of (*β*_*s*_, *β*_*d*_) and (*β*_*s*_, *β*_*d*_, 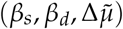), which result in a positive total entropy production and ensure the robustness of the differentiated tissue.

In the retina of a bird, as explained in the previous sections, the five types of photoreceptor cells, namely, green, red, blue, violet, and double cones, form mosaic structures. From the experimental data (see Supporting Information of [60]), we have information about the average standard deviation of the Nearest Neighbor Distribution (NND) for each color as: *σ*_*g*_ ≈ 1.248, *σ*_*r*_ ≈ 1.548, *σ*_*b*_ ≈ 1.729, *σ*_*v*_ ≈ 2.292, and *σ*_*δ*_ ≈ 0.948. In addition, Jiao et al. [61] have found how the variance of an avian cone *σ*^2^ is related to its sensing radius *R* as

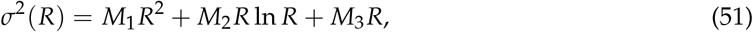

where *M*_1_, *M*_2_, and *M*_3_ have been calculated for each color (see Table I of Ref. [61]). By having this information at our disposal, we are able to calculate the average sensing radius for each photoreceptor as

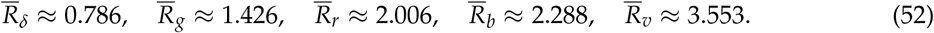

We note that if we assume the minimum sensing radius is equal to the cone size, then the above sensing radii are very close to the average cone size 2.59 ± 1.05*µm* (here, we have used the ±3*σ* rule for the data presented in Fig. 5A of Kram et al. [60]).

In the beginning of this section, we have introduced the variance of the differentiated cells as 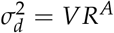; now, by approximating (51) to have such a particular form and setting *V* = 1, and using the values of (52), we can find *A* for each cone as

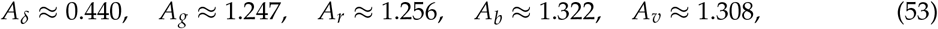

where they are in agreement with the assumption of hyperuniformity made previously, that is, *A* < 2. As an illustration, based on the values of (52) and (53), we have demonstrated the biologically (and physically) acceptable regions of total entropy production (48) in the parametric space of the LEUP parameters: (*β*_*s*_, *β*_*d*_) and (*β*_*s*_, *β*_*d*_, 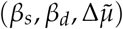) in Fig. 7, in the case of the double cone photoreceptor.

**Figure 7.**
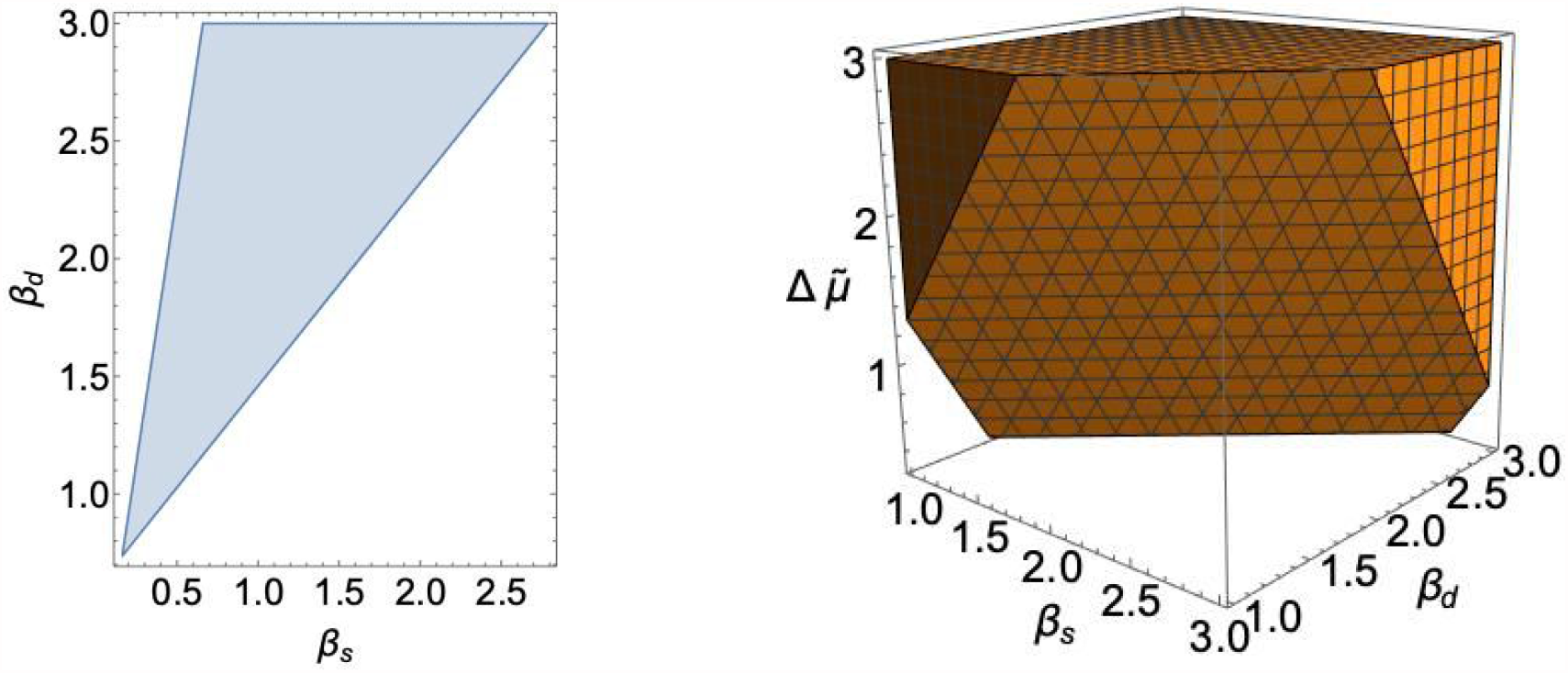
The biologically (and physically) acceptable total entropy production (shaded regions) for the double cone photoreceptor of the avian retina. In the left panel, we have fixed: *C*_0_ = −1, *V* = *U* = 1, *D* = 2, and 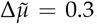. This plot demonstrates the regions where (48) is positive, in the parametric space of the LEUP parameters (*β*_*s*_, *β*_*d*_). We note that we have also imposed the condition regarding the existence of the optimal sensing radius *R*_*c*_, which in this case reads as Γ < 0, see also (49) and (50). In the right panel, we have relaxed the restriction on 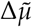.

**Figure 8.**
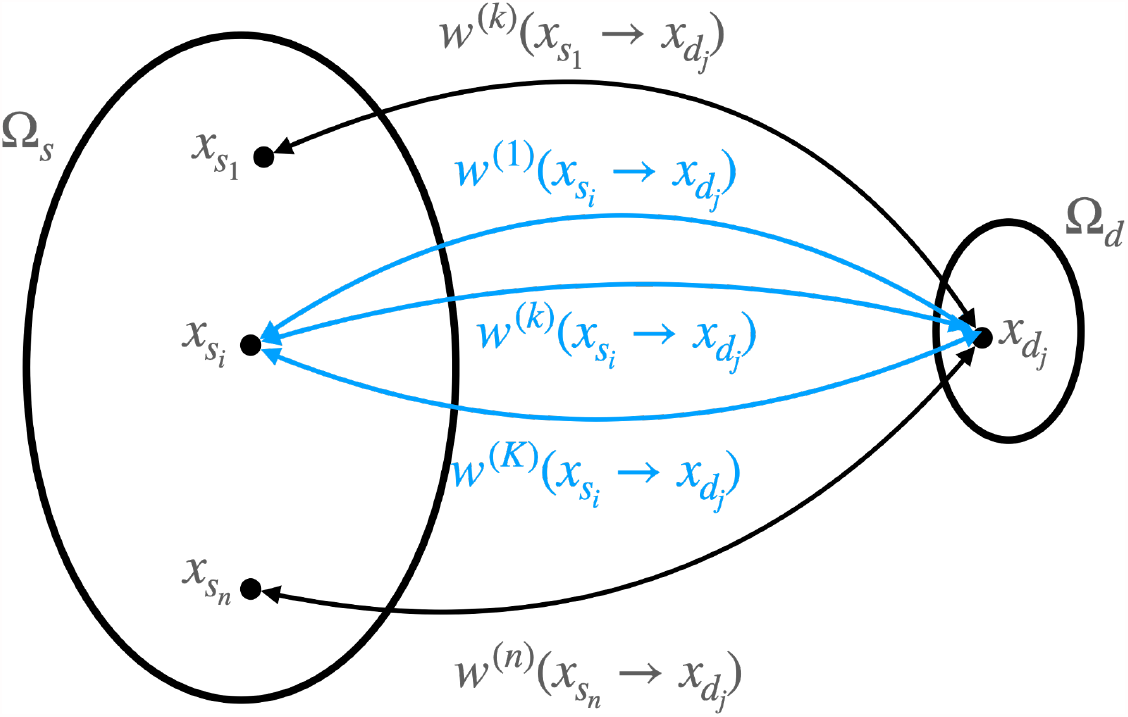
Schematic transitions between pluripotent and differentiated microstates.

At this point, we should remind that the parameter 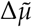 is proportional to the difference of differentiated and pluripotent cells division times. It is well known that the division time of any differentiated cell is much larger than the division time of the corresponding progenitor one [64]. Therefore, it is safe to assume that 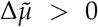. This allows us to identify a relationship for the corresponding *β*’s of the cones and their progenitors, where Fig. 7 implies that *β*_*d*_ > *β*_*s*_. The latter is a sensible result since differentiated cells are expected to be more attentive to their microenvironment in order to ensure optimal cooperation and tissue integrity.

## 6. Discussion

In this paper, we have posed the question **(Q1)** on how cells coordinate intrinsic and extrinsic variables to determine cell decisions that eventually lead to organized and stable tissues. To tackle this problem, we have employed the Least microEnvironmental Uncertainty Principle (LEUP), which has been recently proposed to understand cell decision-making in multicellular systems, and so far it has been applied to cell migration force distribution [25], collective cell migration [26], and binary phenotypic plasticity [27]. In the context of the LEUP, we regard differentiation as a sort of Bayesian decision-making, where cells update their intrinsic variables by encoding microenvironmental information and producing relevant responses. This provides us with a distribution of internal states that depends explicitly on the information of the cell current microenvironment, which is represented by a mesoscopic microenvironmental entropy. Interestingly, we have shown that local microenvironmental entropy should decrease in time leading to more organized cellular microenvironment, which is the case in differentiated tissues. As a proof of principle, we have challenged the LEUP predictions to reproduce differentiated avian photoreceptor mosaics. Although, by fitting a single parameter *β*, we have successfully reproduced the photoreceptor statistics, still this cannot be considered as a rigorous validation. To this end, we have recently gathered an inter-species collection of photoreceptor mosaics to further investigate the potential of the LEUP to reproduce these tissues and possibly classify them.

By using the aforementioned results, we have attempted to shed light on the macroscopic transition between pluripotent and differentiated tissues and have specified it to the formation of photoreceptor mosaics, which is related to the question **(Q2)** posed in Introduction. In this respect, we have developed a stochastic thermodynamic-like theory, based on the Crooks’ theorem, for a general cell and tissue differentiation process. We have shown that differentiated tissues are highly robust to dedifferentiation, even though individual cells are allowed to go back into pluripotent phenotypes. Biologically, the robustness of differentiated tissues depends on reduced proliferation, change from anaerobic to aerobic metabolism, and increased cell sensing that leads to higher order of microenvironmental organization. In particular, we have estimated the critical sensing radii of photoreceptor cones that ensure the thermodynamic robustness of differentiated mosaics, which turns out to be in the range of 0.8*µm* to 3.5*µm*, see (52). We note that the critical radius is the minimal radius required to ensure tissue robustness and therefore should serve as a lower bound for the real values. Now, if we assume that the minimum sensing radius is equal to the cone size, then our predicted range correlates with the average cone size 2.59 ± 1.05*µm* (here, we have used the ±3*σ* rule for the data presented in Fig. 5A of Kram et al. [60]).

In summary, our LEUP-driven model is based on the four crucial assumptions: **(A1)** there is a timescale separation between the internal and microenvironmental variables dynamics, **(A2)** the multicellular system (tissue), where cell is differentiating, follows a Markovian dynamics with the assumption of microscopic reversibility, **(A3)** a flat cell state distribution is assumed for pluripotent cell states, and **(A4)** the spatial distribution of the early microenvironmental pluripotent cells follows a Poisson distribution. Based on these assumptions, we have arrived at three important results: **(I)** predicting the color percentage of the cone cells in the avian retina without any knowledge about the underlying biophysical and biochemical mechanisms, **(II)** demonstrating the robustness of cell-tissue differentiation in thermodynamic terms, and **(III)** determining the limits of the cell sensing radius by establishing a relation between total entropy production and microenvironmental sensing. In the following, we further elaborate on these results.

### Prediction of the cone color distribution

By calibrating a single parameter in the LEUP, we are able to predict the cone color percentage in the avian retina accurately. Our finding regarding the LEUP parameter which reads as *β* ≈ 1.754 (close to 2), gives a strong indication that cells sense their environment quasi-optimally when choosing a particular cell fate during differentiation process. For future study and investigation, we want to examine the validity of this result for photoreceptor mosaics of other species [65–68].

### Robustness and measurement of information gain in differentiation

We have constructed the fluctuation theorem for tissue differentiation and have derived a generalization of the second law of thermodynamics for this process based on a Markovian dynamics. It would be interesting and more realistic to relax this assumption and to analyze the problem using the LEUP on the basis of non-Markovian or memory processes [69], which can lead to different results. The only requirement is that system should have a unique stationary state. We also note that the LEUP-driven second law of thermodynamics can also be seen as a generalization of the Bayesian second law of thermodynamics [70] and the conditional second law of thermodynamics in a strongly coupled system [71].

Our theory, which in the current paper has been applied to the specific case of the avian photoreceptor mosaics, suggests that differentiated tissue is (highly) thermodynamically robust, that is, the arrow of time is almost irreversible, and this robustness depends on microenvironmental sensing and cell metabolism. It should be remarked that, as is demonstrated in (38), if we have the values of 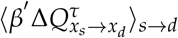 and ⟨Δ*S* ⟩_*s*→*d*_, then we can determine the upper bound of pointwise mutual information difference which is denoted as ⟨Δ*i*⟩ _*s*→*d*_ within the context of the LEUP. However, in the present work, we have only considered transitions between equilibrium end-states and have set ⟨Δ*i*⟩ _*s*→*d*_ = 0. In a future study, we plan to investigate nonequilibrium dynamics of transitions between progenitor and differentiated cell states and establish the upper bound of information gain in cell differentiation. (We also note that the value of ⟨Δ*i*⟩ _*s*→*d*_ can also be obtained directly from experiments, see Refs. [72–75].)

### Limits of sensing radius

By studying total entropy production as a function of cell sensing radius and the LEUP parameters, we have provided an understanding of how a cell regulates its sensing radius according to its microenvironment to ensure the thermodynamic robustness of differentiated tissue. We have shown two cases where **(a)** the entropy production goes to infinity beyond a certain threshold radius which is depicted in Fig. 5 and **(b)** the entropy production goes to a maximum value as in Fig. 6. Interestingly, we conclude that the former is the most biologically relevant case since it requires the division time of differentiated cells to be larger than that of the pluripotent ones and biologically systems operate away from fine-tuned parameter regimes to withstand noisy perturbations. On the technical side, we have assumed that the spatial distribution of pluripotent tissue resembles a Poisson distribution. It would be interesting to relax this assumption and to derive this distribution from real tissue data.

One important issue is the range of validity of **(A1)** regarding the timescale separation between cell decision and cell cycle characteristic times. Although cell decisions may seem happening within one cell cycle, the underlying molecular expressions may evolve over many cell cycles [76,77]. When these molecular expressions cross a threshold then cell decision emerges very fast. Therefore, the definition of cell decision should be treated with care. In our case, we specify a cell decision only when the cell state switches to another dynamic attractor, that induces at the same time some noticeable phenotypic changes. Such attractor transitions are manifested as switches with much shorter characteristic times than a cell cycle [20].

We want to make a brief comment on the relation of our theory to the commonly used approach of the maximum entropy production (MEP). The MEP formalism explains only the transition from a pluripotent state to a differentiating one without realizing the corresponding dynamics. To build a connection between these two, one has to construct the LEUP theory for transition paths like maximum caliber principle [78]. Instead of internal and external variables, one uses internal and external paths of the corresponding evolution. Then, by exploiting the formulation of maximum caliber, one can write the time evolution of microenvironmental path entropy – as a conservation equation – in terms of sources and fluxes and subsequently in terms of path action and entropy production [79,80]. In this regard, one can use maximum caliber principle to construct appropriate transition probabilities and even understand the spatiotemporal dynamics [81]. Finally, one should maximize the internal path entropy which resembles the MEP approach. Working out the details of this connection remains for a future work.

Our proposed theory has important and interesting implications for cancer research and therapy. In particular, (42) states that the balance of metabolic, proliferative, and tissue organization changes (the LEUP term), needed to be taken place in order to destabilize the differentiated state, that is, to promote carcinogenesis. Until now, the majority of the therapies were focused on antiproliferative strategies, such as chemotherapy and radiotherapy, and more seldom to the metabolic conditions such as vasculature normalization. Here, we have proposed that changes in the tissue organization plays a critical role. This fact has been very recently identified in the context of tumor evolution by West et al. [82]. We also note that in [83], it is realized that microenvironment normalization might be the key for immunotherapeutic success. The mechanistic connection between tissue architecture and cell sensing mechanisms is established in the context of our theory. Strikingly, the experimental work of M. Levin’s group [84] shows that disrupting the ion channel sensing in a tissue can induce tumorigenesis. In this regard, we have put forward that investigating changes in the cell sensory processes deserve more attention and might be pivotal in treating cancers. Our goal is to calibrate the existing theory to human photoreceptors data, thus we could apply these ideas to retinoblastoma tumours.

In a nutshell, we have shown how the LEUP facilitates the inference of cellular intrinsic states (or, cell phenotypes) by means of local microenvironmental entropies or fluctuations. This allows the evaluation of cellular states without the detailed grasp of the underlying mechanisms. The sole knowledge about extrinsic variables distributions (or, collective cell decision-making) suffices. Therefore, we can apply the LEUP to cell differentiation problems where the biological or biophysical knowledge is unclear or unknown.

## Author Contributions

All authors contributed equally to this work.

## Appendix: the proof of differentiation microreversibility relation

Let us assume that a particular differentiated state belongs to the corresponding fixed point attractor which involves a number of realizations, 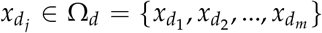. A pluripotent cell state belongs also to an attractor – but not to a fixed point as is discussed in the main text – that involves the following realizations, 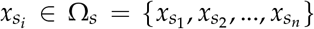, where *n* ≫ *m*. The transition probability between a pluripotent 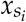 and a differentiated 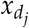 microstate can be denoted as 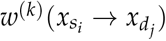, where *k* ∈ {1, …, *K*} represents a possible path between the two states, see also Fig. 8. Now, by invoking the Crooks’ theorem, we can write the condition of microscopic reversibility for a single path *k* as

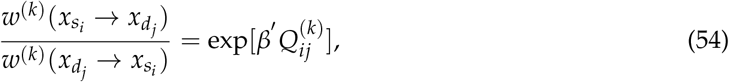

where *β* ′≡ 1/*T*, which *T* is the temperature of heat bath. The quantity 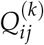 is the heat dissipation in the *k*-th path during the transition from 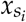 to 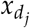. Now, by averaging over all paths, we can write the path-independent transition probability as

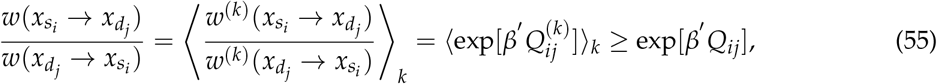

where 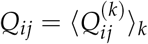 is the total heat dissipation over all paths (the lower bound is based on Jensen’s inequality). The transition between a pluripotent state, *x*_*s*_, to any differentiated state, *x*_*d*_, can be interpreted as a transition from any 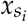 to any 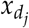. Therefore, we can write:

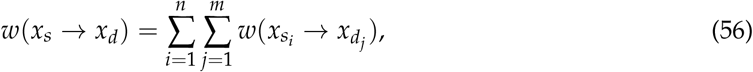

where the average over the corresponding paths is assumed. Now, the ratio of the forward/differentiation over the backward/dedifferentiation transition probability reads

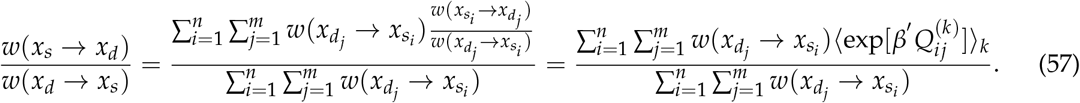

The last term in (57) can be viewed as a weighted average over all possible dedifferentiation paths between pluripotent and differentiated states, that is,

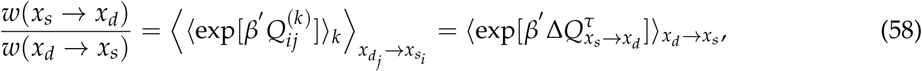

where the latter has been used for notational simplicity. This is the microreversibility relation for a general differentiation process that corresponds to (22) in the main text.

